# Classification of *Wolbachia* (Alphaproteobacteria, Rickettsiales): No evidence for a distinct supergroup in cave spiders

**DOI:** 10.1101/046169

**Authors:** Michael Gerth

**Author notes:** Contact Michael Gerth Institute for Integrative Biology University of Liverpool Biosciences Building Liverpool L69 7ZB United Kingdom.

## To the editor

*Wolbachia* are intracellular, inherited Alphaproteobacteria present in a large proportion of arthropod species and in filarial nematodes. *Wolbachia–host* interactions are many-faceted and include, but are not limited to reproductive manipulations (Werren et al. 2008), nutritional mutualism (Nikoh et al. 2014), and protection from pathogens (Hedges et al. 2008, Teixeira et al. 2008). Although there is just a single *Wolbachia* species described, the genus is highly diverse with regard to its distribution, phenotypes induced in the host, and genomic architecture. The generally accepted classification scheme currently in use differentiates among genetically distinct, monophyletic lineages named “supergroups” (Lindsey et al. 2016). To date, 16 of these *Wolbachia* lineages that – based on multiple genetic markers – are clearly distinct from another were described (supergroups A–F and H–Q, Glowska et al. 2015).

In a recent study, Wang et al. (2016, from here on "Wang et al.") reported the discovery of a novel, 17^th^ *Wolbachia* supergroup (“R”) from cave spiders *(Telema* ssp.). The authors base this conclusion on phylogenetic analyses of sequences from three protein coding genes *(ftsZ, coxA, groEL*) and the 16S rRNA gene. Here, I re-analyse these data and show that *Wolbachia* from *Telema* spiders clusters with supergroup A strains and thus, the creation of a novel *Wolbachia* supergroup for these strains is without justification.

Using the sequence data generated by Wang et al., I first performed an online BLAST search of all *ftsZ, coxA, groEL*, and *16S* sequences (2, 6, 1, and 5 haplotypes, respectively) against the nucleotide database of NCBI GenBank. Strikingly, all of the sequences match to known supergroup A strains with an identity greater than 99%. Next, using Mafft version 7.245 (Katoh and Standley 2013) with automatic parameter settings, I aligned all novel sequences with the corresponding orthologs extracted from the *Wolbachia* genomes wRi, wHa, wMel (all supergroup A), wPip & wNo (supergroup B), and wCle (supergroup F). Using Aliview version 1.17.1 (Larsson 2014), the datasets were then trimmed manually to include only positions that were sequenced by Wang et al.

All alignments are available under https://github.com/gerthmicha/supergroup_R. From these alignments, raw genetic distances were calculated using the “ape” package within the R statistical environment (Paradis et al. 2004, R Core Team 2016). All *Wolbachia* loci from *Telema* hosts are very similar to each other and to the reference supergroup A loci (~0-1% distance, Fig. S1). In general, the distances within supergroup A (including *Telema* strains) are almost one order of magnitude smaller compared to the distances between supergroups (Fig. S1). The distance measures are therefore in line with the conclusion that *Wolbachia* from cave spiders are supergroup A strains.

Finally, maximum likelihood trees were created for all single gene alignments and a supermatrix of all concatenated genes using IQ-TREE multicore version 1.4.1 (Nguyen et al. 2015) with automated model selection (“-m TEST”) and 1000 ultrafast bootstrap replicates. All of the single gene trees and the tree based on the concatenated genes strongly support an association of *Wolbachia* strains from *Telema* ssp. with supergroup A (Fig. 1). This becomes especially clear when comparing the genetic distances within supergroup A and the distances between supergroups A, B, and F (Fig. 1).

**Fig. 1.**
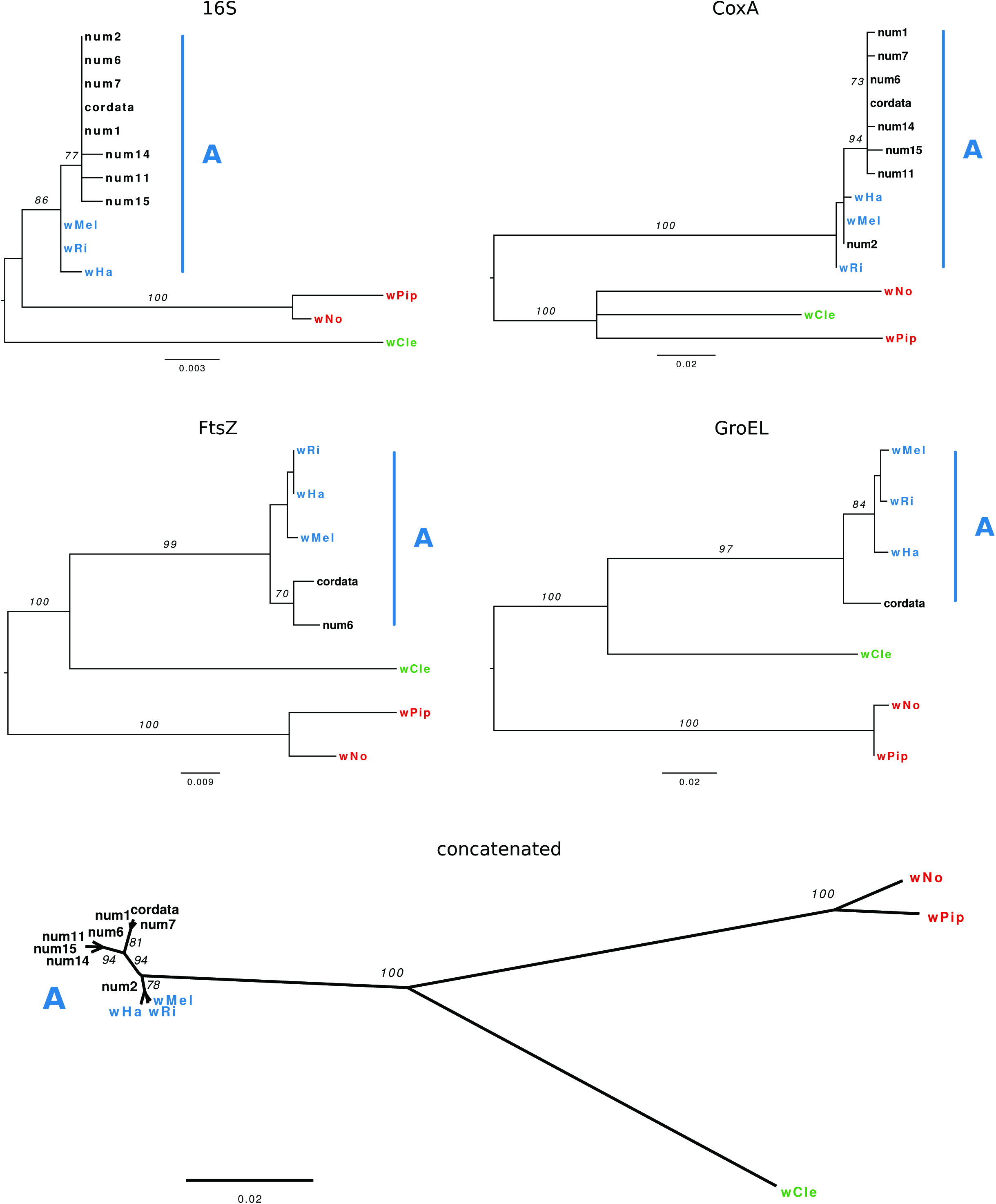
Phylogenetic analyses of *Wolbachia* sequences isolated from *Telema* ssp. (cave spiders). Maximum likelihood trees are shown for each single gene analysis and for the concatenated alignment of all four genes. Numbers on branches correspond to bootstrap values from 1000 pseudoreplicates. Only values above or equal to 70 are shown. Single gene trees are midpoint rooted. Reference sequences are highlighted as follows: blue-supergroup A, red-supergroup B, green-supergroup F. Naming of the sequences from *Telema* hosts follows Wang et al. (2016).

Visual comparisons of the trees presented in Fig. 1 and the ones generated by Wang et al. reveal little discordance. Single gene trees from maximum likelihood analyses of *coxA, ftsZ*, and *16S* sequences (Supplementary Figure 1 in Wang et al.) show a very close association of *Wolbachia* sequences from *Telema* ssp. with supergroup A that was also recovered in the re-analyses presented here (although for *16S*, no supergroup A reference sequences were included by Wang et al.). However, the analysis of *groEL* sequences (Supplementary Figure 1D in Wang et al.) and the one of the concatenated dataset resulted in a placement of *Wolbachia* from *Telema* ssp. distinct from other supergroups. From the data available, these analyses are not replicable. Speculatively, an alignment artefact has led to the erroneous placement of *Wolbachia* from *Telema* ssp. in the analysis of Wang et al.

The discordance between single gene trees and the concatenated analysis was noted by

Wang et al. and led them to conclude that a single locus is not sufficient to resolve phylogenetic relationships within *Wolbachia*. Whilst this is certainly true, the observation that each single gene analysis consistently places *Wolbachia* from *Telema* ssp. within (or closely associated to) supergroup A questions the validity of the supermatrix analysis. Further to this, the fact that *Wolbachia* from *Telema* ssp. form a monophyletic group in most of the analyses does not imply that they should be considered a separate supergroup, as Wang et al. suggest. Naturally, each *Wolbachia* supergroup is composed of multiple monophyletic lineages. The reference sequences used here and by Wang et al. do not reflect the full genetic diversity of supergroup A strains (which are very common among arthropods), and while *Wolbachia* from *Telema* ssp. may seem distinct in these analyses, this is not necessarily the case when considering the multitude of supergroup A alleles of the loci in question. Finally, the detection of supergroup A-like prophage sequences in *Telema* hosts provides further evidence for their inclusion in this supergroup.

In summary, the here presented similarity searches, genetic distance measures, and phylogenetic analyses are all consistent and provide strong evidence for the association of *Wolbachia* strains from *Telema* with supergroup A. Their placement within a novel supergroup, as proposed by Wang et al., lacks any support and is likely artefactual. In order to minimize confusion, it may be helpful not to assign the name “supergroup R” to any novel *Wolbachia* lineage potentially to be discovered in the future.

## Acknowledgements

I am grateful to Guan-Hong Wang for providing sequence data and for discussions. I thank Franziska Anni Franke for comments on the manuscript. My current position is funded by EMBO (ALTF 48-2015) and co-funded by Marie-Curie Actions of the European Commission (LTFCOFUND2013, GA-2013-609409).

## Supplementary information

**Fig. S1.**
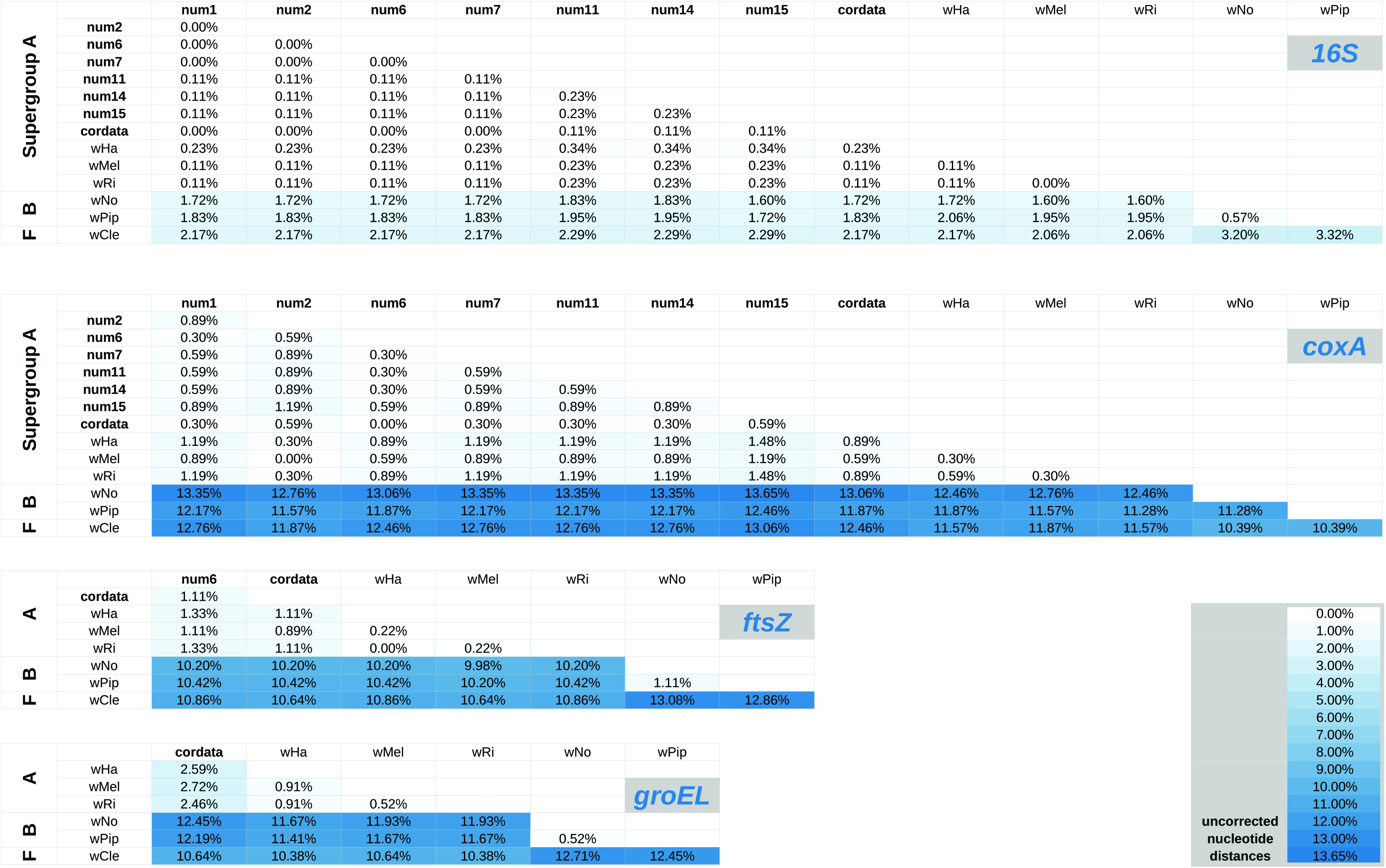
Distance matrices (uncorrected p-distances) for each alignment analysed in this study. Sequences from *Telema* sp. are highlighted in bold.

## References

Glowska, E., Dragun-Damian, A., Dabert, M. & Gerth, M. (2015). New Wolbachia supergroups detected in quill mites (Acari: Syringophilidae). Infect. Genet. Evol, 30, 140–146.

Hedges, L. M., Brownlie, J. C., O'Neill, S. L. & Johnson, K. N. (2008). Wolbachia and virus protection in insects. Science, 322, 702.

Katoh, K. & Standley, D. M. (2013). MAFFT multiple sequence alignment software version 7: improvements in performance and usability. Mol. Biol. Evol, 30, 772–780.

Larsson, A. (2014). AliView: a fast and lightweight alignment viewer and editor for large datasets. Bioinformatics, 30, 3276–3278.

Lindsey, A. R., Bordenstein, S. R., Newton, I. L. & Rasgon, J. L. (2016). Wolbachia pipientis should not be split into multiple species: A response to Ramirez-Puebla et al.,“Species in Wolbachia? Proposal for the designation of ‘Candidatus Wolbachia bourtzisii’,‘Candidatus Wolbachia onchocercicola’,‘Candidatus Wolbachia blaxteri’,‘Candidatus Wolbachia brugii’,‘Candidatus Wolbachia taylori’,‘Candidatus Wolbachia collembolicola’ and ‘Candidatus Wolbachia multihospitum’for the different species within Wolbachia supergroups”. Syst. Appl. Microbiol, doi:10.1016/j.syapm.2016.03.001.

Nguyen, L.-T., Schmidt, H. A., von Haeseler, A. & Minh, B. Q. (2015). IQ-TREE: A fast and effective stochastic algorithm for estimating maximum-likelihood phylogenies. Mol. Biol. Evol, 32, 268–274.

Nikoh, N., Hosokawa, T., Moriyama, M., Oshima, K., Hattori, M. & Fukatsu, T. (2014). Evolutionary origin of insect-Wolbachia nutritional mutualism. Proc. Natl. Acad. Sci. U.S.A, 111, 10257–10262.

Paradis, E., Claude, J. & Strimmer, K. (2004). APE: analyses of phylogenetics and evolution in R language. Bioinformatics, 20, 289–290.

R Core Team (2016). R: A language and environment for statistical computing. R Foundation for Statistical Computing, Vienna, Austria.

Teixeira, L., Ferreira, A. & Ashburner, M. (2008). The bacterial symbiont Wolbachia induces resistance to RNA viral infections in Drosophila melanogaster. PLoS Biol, 6, e1000002.

Wang, G.-H., Jia, L.-Y., Xiao, J.-H. & Huang, D.-W. (2016). Discovery of a new Wolbachia supergroup in cave spider species and the lateral transfer of phage WO among distant hosts. Infect. Genet. Evol, doi:10.1016/j.meegid.2016.03.015.

Werren, J. H., Baldo, L. & Clark, M. E. (2008). Wolbachia: master manipulators of invertebrate biology. Nat. Rev. Microbiol, 6, 741–751.

